# Diverse rhizosphere-associated *Pseudomonas* genomes isolated along the marine-terrestrial transition zone of a Wadden island salt march

**DOI:** 10.1101/2023.11.14.566819

**Authors:** Siyu Mei, Miao Wang, Joana Falcão Salles, Thomas Hackl

## Abstract

Soil microbes are key drivers of ecosystem processes promoting nutrient cycling, system productivity, and resilience. While much is known about the roles of microbes in established systems, their impact on soil development and the successional transformation over time remains poorly understood. Here, we provide 67 diverse, rhizosphere-associated *Pseudomonas* draft genomes from an undisturbed salt march primary succession spanning >100 years of soil development. *Pseudomonas* are cosmopolitan bacteria with a significant role in plant establishment and growth. We obtained isolates associated with *Limonium vulgare* and *Artemisia maritima*, two typical salt marsh perennial plants with roles in soil stabilization, salinity regulation, and biodiversity support. We anticipate that our data, in combination with the provided physiochemical measurements, will help identify genomic signatures associated with the different selective regimes along the successional stages, such as varying soil complexity, texture, and nutrient availability. Such findings would not only advance our understanding of *Pseudomonas’* role in natural soil ecosystems but also provide the basis for a better understanding of the roles of microbes throughout ecosystem transformations.

## Background & Summary

Advancements in cell culture, DNA extraction, and sequencing technologies have greatly increased our understanding of the variable complexity and diversity of soil microbial communities. Latest technologies enable the collection of samples *in situ* and provide information on the gene level of the soil microorganisms, which can then be used to infer their ecological roles. One aspect that has interested microbial ecologists is understanding the genomic modifications that allow soil microorganisms to colonize soils during soil development. However, the ability to sample over time spans necessary for soil formation is challenging^1^. Primary succession or chronosequence have been identified as valuable playgrounds to address these questions due to their associated space-for-time substitution. As such, they have been used to study soil development and functional changes related to the variation in above-ground and below-ground processes^2,3,4^ providing critical information over temporal community dynamics and soil development across multiple time scales^5,6^ From a microbial perspective, several researchers have utilized chronosequences to study soil communities in different areas, such as grassland^3^, forests^4^, deglaciated soils^5^, and salt marshes^7^.

Salt marshes are tidal wetlands that play a vital ecological role in the coastal ecosystem and maintain water quality and habitat health by filtering pollutants, runoff, and excess nutrients. Moreover, they store carbon at a rate ten times that of mature tropical forests, helping moderate climate change effects^8^. The salt marsh located on the Waddensea barrier island of Schiermonnikoog constitutes a well-documented chronosequence covering more than one hundred years of succession^9^. Recent studies revealed that soil microbial communities change in taxonomic composition throughout this chronosequence, with soil organic matter and salt concentrations being the major drivers of soil microbial community structure^10^. These soil microbial communities also differ functionally and are enriched in genes associated with dispersal at the early stages. In contrast, the late stages exhibit enrichment in antibiotic resistance genes^5^. While these studies have provided important insights into the effects of succession at the community level, inter- and intraspecies variation remains poorly understood.

*Pseudomonas* is one of the most studied and diverse bacterial genera because of its prevalence in several environments, such as soil^11^ and water^12^, and macroscopic hosts, such as plants^13^, mammals^14^, and insects^15^. *Pseudomonas* is ecologically well-studied in soil under anthropogenic influence and is regarded as a key indicator species to reflect disturbances. The diversity and structure of *Pseudomonas* communities significantly correlate with the changes in soil fertilization^16^ and the long-term use of mineral fertilizers^17^. Moreover, the genus *Pseudomonas* contains several plant-associated and free-living beneficial strains commonly found in soils from sustainable crop production, where they can act as plant growth-promoting or biocontrol agents^18^. However, research thus far has focused on cultivated soils, while *Pseudomonas* isolates from lesser explored niches show beneficial properties and genetic heterogeneity^19^, clearly indicating an untapped potential.

Here, we used the genus *Pseudomonas* as a focus group to determine how the changes in soil characteristics along the primary succession of Schiermonnikoog lead to changes in their genomes. For that, we isolated *Pseudomonas* strains from the endo- and rhizosphere of typical salt marsh plants with broad distribution along the chronosequence^20^. We provide whole genome sequencing data of 67 *Pseudomonas* and the physiochemical parameters from the soil samples collected from the roots of salt marsh plants on the island of Schiermonnikoog, the Netherlands. These data represented snapshots of the *Pseudomonas* community spanning a time scale of 100 years, with successional ages ranging from 5 to 105 years (Figure 2). From each location, we sampled rhizosphere soil and root endosphere from two plants, *Limonium vulgare* and *Artemisia maritima*. The complete data contains the raw sequencing reads, the cleaned assemblies with the taxonomic information collected through sampling, measurement, cultivation of individual isolates, DNA extraction, whole genome sequencing, and the subsequent refinement and annotation steps (Figure 1). We expect that these data will help study microbial questions from the perspective of gene level, including, but not limited to, microbiology, microbial ecology, genetics, and evolution. In addition, the associated physiochemical parameters, in particular, might link the microbial world and environmental disturbances.

**Figure 1:**
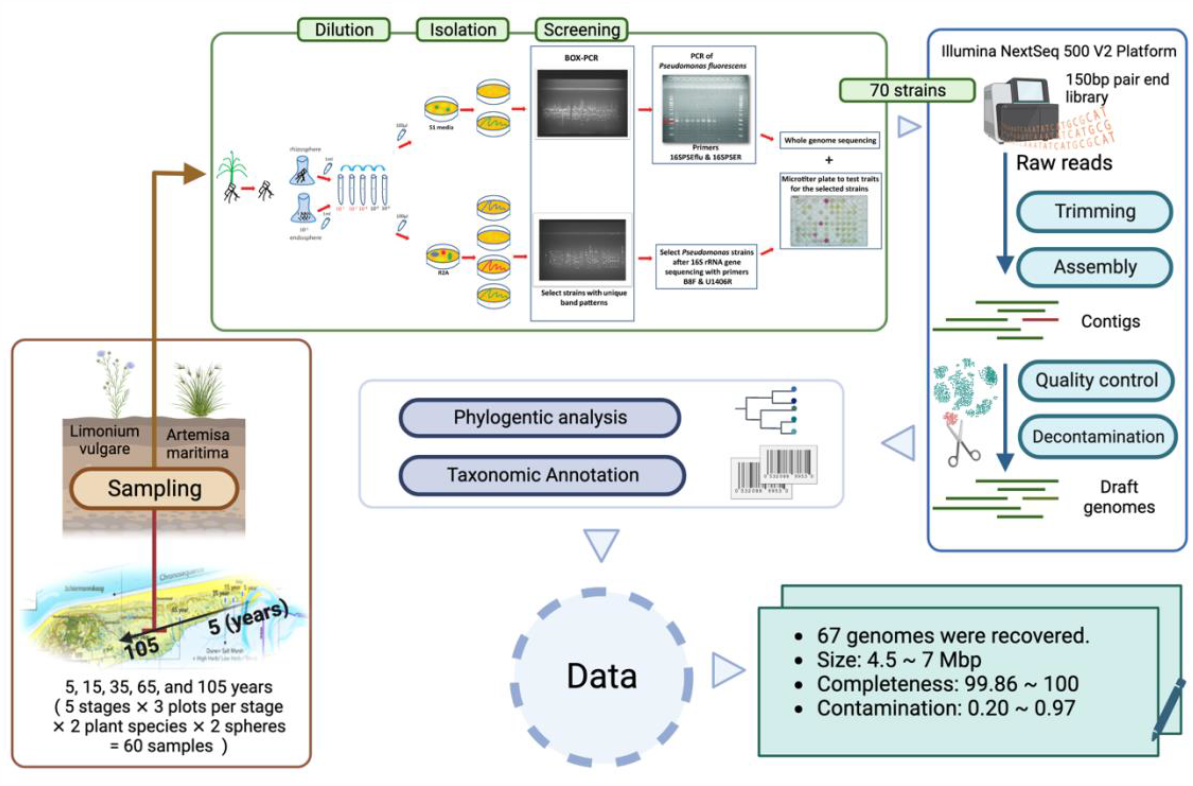
Overview of sampling procedure, sample processing, genome sequencing, and bioinformatic analysis. Diagram illustrating the main stages and procedures for generating the dataset of this study: endo- and rhizosphere sampling, strain isolation, whole-genome sequencing, and assembly and bioinformatic analysis. Summary metrics of all genomes are provided in the bottom-left box. The figure was created with BioRender.com
.

**Figure 2:**
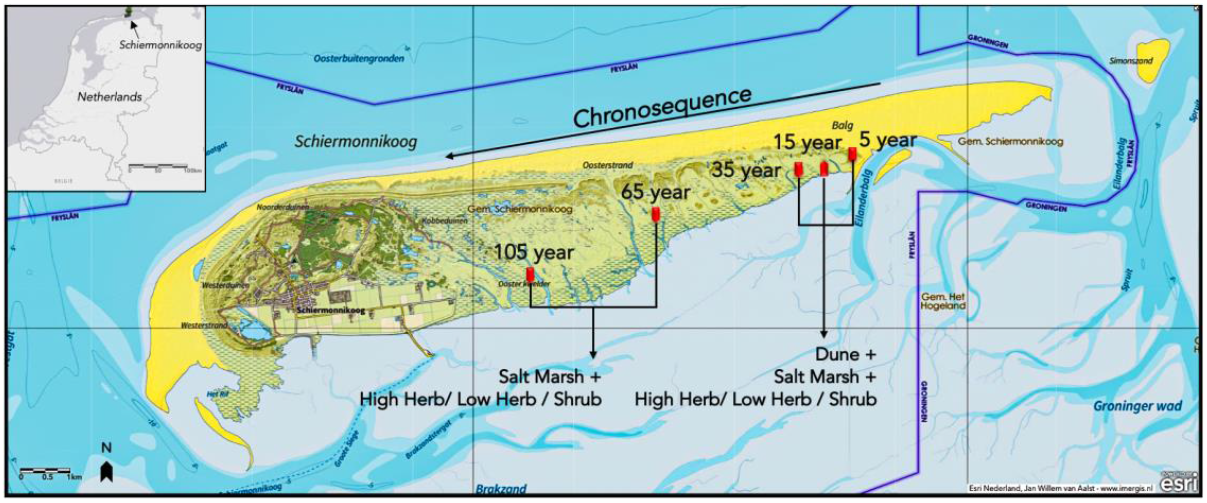
Sampling locations along the chronosequence on the island of Schiermonnikoog. Map of Schiermonnikoog displaying the location where whole genomes were collected from the indicated soil types. Wadden islands like Schiermonnikoog constantly migrate, with soil washed away from the West end and new sand continuously deposited at the East end. Our sampling locations (red bold arrows) are part of a well-documented chronosequence representing more than 100 years of soil development (long dark arrow). The soil stages by years of soil formation are indicated above the location symbols. The type of vegetation habitats is indicated by dark arrows and text below the symbols of locations. This Figure was completed on ArcGis Online, and the base map was obtained from Esri Nederland^37^.

## Methods

Plants and root-attached soil were collected in April 2016 from five sites along a successional chronosequence on the Dutch Wadden island Schiermannikoog (soil ages of 5, 15, 35, 65, and 105 years)^20^. Within each plot, four healthy-looking *L. vulgare* and *A. maritima* of similar sizes with soil adhering to the intact roots were obtained and processed together, generating two composite samples per plot. Thirty composite samples in total were collected (5 stages ⨯ 3 plots per stage ⨯ 2 plant species). Each sample was placed in a sterile plastic bag, sealed, and transported to the laboratory within 24 hours. From each composite sample, we sampled rhizosphere soil and root endosphere. For rhizosphere sample, 8-10 g of soil was diluted in 47ml 1 x PBS solution, shaken for 30 min at 200 rpm at room temperature, and prepared for serial dilutions (1/10) in sterile 1X PBS. For endosphere samples, the roots were washed under running water, trimmed to remove adhering soil and dead tissues, and surface-sterilized^21^. The samples were then diced with a sterile scalpel and immersed into 45 ml 0.9% NaCl solution. After incubation for 1h at 28 °C, the suspension was shaken using a horizontal vortex instrument, followed by serial dilution (1/10) in sterile 1X PBS. Sterility checks were performed by tissue-blotting surface-sterilized root samples on R2A plates at 28 °C for 2–7 days. Only samples without bacterial growth were considered successfully sterilized and used further.

*Pseudomonas* isolates were obtained using Gould’s S1 medium, a selective medium previously used for isolating fluorescent *Pseudomonas*^22^. Thirty-two bacterial colonies per plate with unique morphologies were purified using a streak-plate procedure, transferred onto new S1 and R2A^23^ medium plates, and further used as templates for BOX-PCR, which is a DNA-based typing method capable of simultaneously screening many DNA regions scattered in the bacterial genome^24^. 109 bacterial cultures with unique BOX-PCR patterns on Gould’s S1 agar plates were subjected to total DNA extraction using the MoBio UltraClean Microbial DNA Isolation Kit (MoBio Laboratories, Carlsbad, CA, USA). All DNA samples were standardized to an equal concentration of 5 ng μL^-1^ for sequencing. 70 *Pseudomonas* spp unique isolates were sent to LGC Genomics GmbH (Berlin, Germany) for whole genome sequencing on an Illumina NextSeq 500 V2 platform with a 150-bp paired-end library design.

We trimmed the resulting Illumina reads by using Trim Galore^25^, then assembled the trimmed reads de novo with SPAdes^26^, aiming for genome sizes around 6 Mbp with GC contents of 60%-70%^27^ (*Pseudomonas aeruginosa* 65–67%, size 5.5–7 Mbp; *Pseudomonas fluorescens∼*60%, size∼6 Mbp^28^). We kept only scaffolds larger than 1000 bp (Table S1) and further decontaminated the raw contigs to obtain high-quality draft genomes (see Technical Validation for more information). We performed taxonomic classification using the Genome Taxonomy Database (GTDB) and the associated toolkit (GTDBtk)^29^. Detailed taxonomic information is provided in Table S4. To further assess the evolutionary relationships among the 67 genomes of this study, we reconstructed a phylogenetic tree (Figure 4). We used 34 reference genomes, including 9 outgroup species^30^ and the 25 closest reference genomes identified by GTDBtk (NCBI genome accession listed in Table S5). We used GTDBtk to extract 120 universal bacterial marker amino acid alignments from all genomes and FastTree^31^ to reconstruct a phylogenetic tree based on the amino acid sequence alignments. The resulting tree was visualized with ggtree^32^ and Evolview^33^.

## Data Records

The raw Illumina sequencing reads are available from the European Nucleotide Archive (ENA). Accession numbers, library size, and coverage statistics can be found in Table S2. The curated and annotated assemblies have been deposited in <insert link>. Physiochemical parameters can be found in Table 1.

**Table 1:**
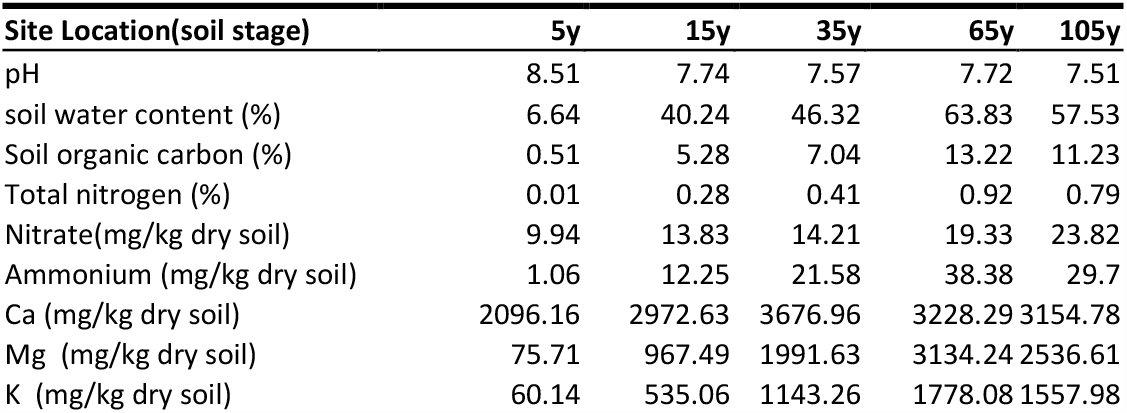
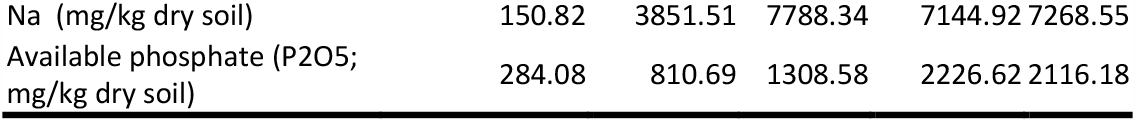
Physiochemical parameters of soil along the sampling locations.

## Technical Validation

The overall sequencing quality of the short-read data was assessed with FastQC^34^ and low-quality regions were trimmed with Trim Galore^35^. To ensure high-quality genomes, assemblies were manually screened for low-coverage contamination (Figure S1), and suspicious contigs were removed. In total, we recovered 67 genomes with good quality based on CheckM^36^ metrics (Completeness > 80%, contamination > 10%, completeness – 5*contamination > 80%) (Figure 3).

**Figure 3:**
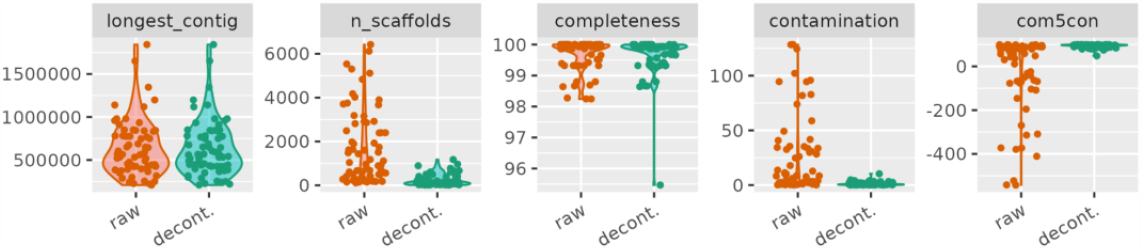
*In-silico* decontamination considerably improves *Pseudomonas* genome assembly quality. Quality metrics of genome assemblies before (orange) and after (green) decontamination: the size of the longest contig, number of scaffolds, completeness, contamination, and integrated score calculated as completeness - 5*contamination. All values were computed with CheckM^23^. Quality improvements indicate that manual decontamination based on sequence coverage and length (Figure S1) removed contaminations efficiently without affecting assembly completeness.

**Figure 4:**
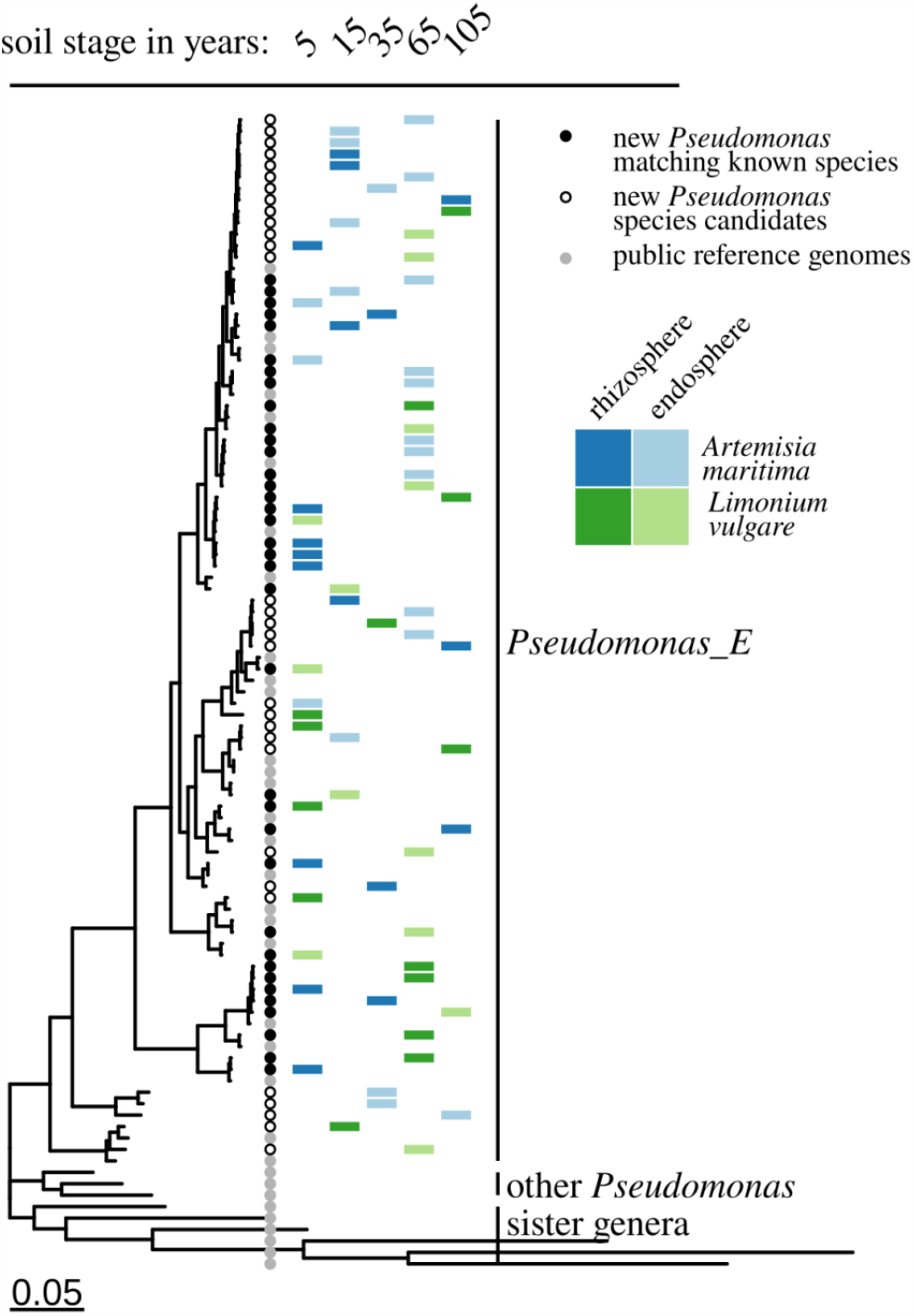
Phylogenetic relationships of 67 *Pseudomonas* isolates and their distribution across soil stages, host plants, and habitat. Phylogenetic tree including nine outgroup species (first 9 species from the bottom), *26 Pseudomonas_E* reference genomes, and the 67 *Pseudomonas* genomes of this study. New genomes were assigned to known species based on closest placement in the Genome Taxonomy Database (GTDB)^28^. 31 genomes did not match at species-level cutoff (ANI >=95 %, AF>=0.5) to known *Pseudomonas* and can be considered candidates for new species. Isolates were obtained from 5 different soil stages (5 years to 105 years), from two host plants (*Artemisia maritima* and *Limonium vulgare*) and from both endo- and rhizosphere.

## Figures

All the figures can also be found in the attachments.

## Figure Legends

**Figure S1: Contamination assessment of contigs by coverage, GC content and sequence length**. Each dot represents a contig in the respective assembly with coverage on the y-axis, GC content on the x-axis, and dot size corresponding to the contig length. The horizontal bar indicates the coverage cutoff which we applied to remove contaminations.

## Tables

**Table S1: Genome metadata and raw assembly statistics.**

**Table S2: Reference genomes used for taxonomic analyses**.

**Table S3: Genome assembly metrics before and after decontamination.**

**Table S4: Taxonomic classification summary generated with GTDBtk**.

Table S5: The accession number (waiting for ENA to generate)

## Code availability

Software versions and any relevant variables and parameters employed are as follows:

- Trim Galore v0.6.5 (trim_galore --paired --fastqc --phred33 --illumina)^25^;
- FastQC v0.11.9^34^;
- SPAdes v3.15.4 (spades.py --careful -t 20)^26^;
- Seqkit v2.3.0 (seqkit grep -f)^38^;
- CheckM v1.1.3-foss-2021a (checkm lineage_wf -x fa)^36^;
- GTDB v 1.7.0 (gtdbtk classify_wf); GTDB-Tk reference data version r202^29^;
- FastTree v2.1.11^31^;
- ggtree v3.9.1^32^.

## Acknowledgments

We thank Han Olff, Matty Berg, Chris Smit, Maarten Schrama, and Ruth Howison for information on sampling locations and plant species. We are grateful to Jolanda K. Brons and Armando Cavalcante Franco Dias for sampling expeditions. We thank the ‘Nederlandse Vereniging voor Natuurmonumenten’ for granting us access to the salt marsh. We thank the Center for Information Technology of the University of Groningen for their support and for providing access to the Peregrine and Hábrók high-performance computing cluster. We thank the China Scholarship Council (CSC), on a personal grant to SM (Grant No. [2020]596) and MW (Grant No. [2013]3009).

## Author contributions

MW and JFS designed the study. MW carried out the field and lab work. SM, JFH and TH designed the data analysis. SM carried out the data processing and analysis with contributions from TH. SM and TH wrote the manuscript with contributions from JFS. TH and JFH supervised the project.

## Competing interests

The authors declare no competing interests.

## Notes

### Competing Interest Statement

The authors have declared no competing interest.

